# Hair regeneration by small molecules that activate autophagy

**DOI:** 10.1101/303495

**Authors:** Min Chai, Meisheng Jiang, Laurent Vergnes, Xudong Fu, Stéphanie C. de Barros, Jing Jiao, Harvey R. Herschman, Gay M. Crooks, Karen Reue, Jing Huang

## Abstract

Hair plays important roles, ranging from the conservation of body heat to the preservation of psychological well-being. Hair loss or alopecia affects millions worldwide and can occur because of aging, hormonal dysfunction, autoimmunity, or as a side effect of cancer treatment (Gilhar et al., 2012; Petukhova et al., 2010). Methods that can be used to regrow hair are highly sought after, but lacking. Here we report that hair regeneration can be stimulated by small molecules that activate autophagy, including the longevity metabolites α-ketoglutarate and α-ketobutyrate, and the prescription drugs rapamycin and metformin which impinge on TOR and AMPK signaling.

## Introduction

The biological and psychological importance of hair is well recognized. Mammalian hair growth consists of cyclic repetitions of telogen (quiescence), anagen (regeneration) and catagen (degeneration) phases of the hair follicle (Muller-Rover et al., 2001; Schneider et al., 2009). This hair follicle cycle is regulated by both intrinsic and extrinsic signals which control quiescence and activation of hair follicle stem cells (HFSC). Inadequate HFSC activation and proliferation underlie alopecia in numerous biological and pathological conditions, including aging (Keyes et al., 2013; Chueh et al., 2013). Molecules that can promote HFSC activation and anagen initiation are intensely searched for, as they may help reveal how hair regeneration is regulated and provide therapeutic and cosmetic interventions. Here, we postulate that telogen hair follicles may be induced to enter anagen by pharmacologically triggering autophagy.

As a fundamental process for degrading and recycling cellular components, autophagy is critical for the adaptation to nutrient starvation and other adverse environmental conditions (Galluzzi et al., 2014). Autophagy is also important for quality control of proteostasis through the elimination of misfolded or damaged proteins and damaged organelles. The loss of autophagy may be causally related to neurodegeneration and other diseases (Mizushima et al., 2008). Autophagy declines with age (Levine and Kroemer, 2008), likely accounting for the higher prevalence of autophagy-related diseases (e.g., cancer and neurodegenerative diseases) in the elderly. Autophagic clearing of active, healthy mitochondria in hematopoietic stem cells is required to maintain quiescence and stemness (Ho et al., 2017), and autophagy fulfills the nutrient demand of quiescent muscle stem cell activation (Tang and Rando, 2014). In the skin, autophagy is required for self-renewal and differentiation of epidermal and dermal stem cells (Salemi et al., 2012), but its role in hair follicle stem cells has remained controversial. On one hand, autophagy may be required for hair growth as skin grafts from the autophagy-related gene 7 (Atg7)-deficient mice exhibit abnormal hair growth (Yoshihara et al., 2015). On the other hand, psychological stress induces autophagy and delay of hair cycle (Wang et al., 2015).

Previously, alterations in intrinsic signaling, gene expression, and circadian function were implicated to prevent anagen entry in aged HFSC and result in alopecia (Castilho et al., 2009; Keyes et al., 2013; Solanas et al., 2017). The unforeseen finding that supplementation of a metabolite α-ketobutyrate (α-KB) in old mice can increase longevity and prevent alopecia (Huang et al., 2016) suggests that rejuvenating aging or aging associated deficiencies may restore HFSC function and hair growth in skin tissue. We report herein that autophagy is increased during anagen phase of the natural hair follicle cycle and that autophagy induction by specific small molecules can be used to promote hair regeneration.

## Results

Induction of autophagy is mediated by some of the same cellular energy metabolism regulators which have been linked to or implicated in the effect of dietary restriction (DR) on longevity. We previously showed that the longevity metabolite α-ketoglutarate (α-KG) is induced upon DR and mediates its longevity effect in *C. elegans*. α-KG also increases autophagy in both worms and cultured mammalian cells (Chin et al., 2014). Here we tested whether α-KG can increase hair regeneration using an established in vivo C57BL/6J mouse dorsal skin model. Male mice at 6.5 weeks of age (postnatal day 44) were shaved on the back, when dorsal skin hair follicles are in telogen. α-KG or vehicle control treatment was applied topically every other day. As shown in Figure 1A, α-KG treatment drastically enhances hair regeneration. Since inflammation and wound repair are known to stimulate tissue, including hair, regeneration (Ito et al., 2007), our study only focused on molecules that do not cause skin damage or other abnormal skin conditions. There was no evidence of skin irritation or inflammation by α-KG or other small molecule treatment described in this study (see below). In telogen skin, α-KG initiated new anagen waves as early as 7 days post-treatment. Anagen in black mice is macroscopically recognizable by the melanin pigment visible through the skin, as the melanogenic activity of follicular melanocytes is strictly coupled to the anagen stage of the hair cycle (Slominski and Paus, 1993). In the experiment shown in Figure 1A, skin pigmentation was visible by day 12 post-treatment with α-KG (Figure 1B). In contrast, in vehicle-treated control mice no pigmentation or only a few scattered pigmented spots were apparent at least until day 39, when animals were sacrificed for histological and biochemical analyses. Hair grew from the pigmented skin area of α-KG treated mice within 5~7 days, and by day 39 post-treatment α-KG treated mice exhibited robust hair growth while control mice showed no hair growth overall (Figure 1A). The effects of α-KG on anagen initiation and hair regeneration were even more dramatic when mice were treated later in telogen at 8 weeks of age (Figure S1A-B). α-KG also exhibits similar hair stimulating effect in female mice (Figure S1C). Formation and differentiation of hair follicles in α-KG treated mice were also demonstrated by histological analyses (Figure 1C). More follicles were observed after α-KG treatment, showing anagen phase induction. Further, mice of the same age as those used for regeneration experiments were acutely treated for 5 days and analyzed for early biochemical changes. Increased autophagy induction in the α-KG treated mouse skin was supported by western blot analysis of LC3 expression (Figure 1D). Expression of the autophagy substrate p62/SQSTM1, which is widely used as an indicator of autophagic degradation, was also increased with autophagy induction by α-KG in the mouse skin (Figure 1D) as well as by rapamycin induced autophagy (see below). This is likely due to compensation through upregulation of p62 transcription, as was reported for p62 during prolonged starvation (Sahani et al., 2014).

**Figure 1.**
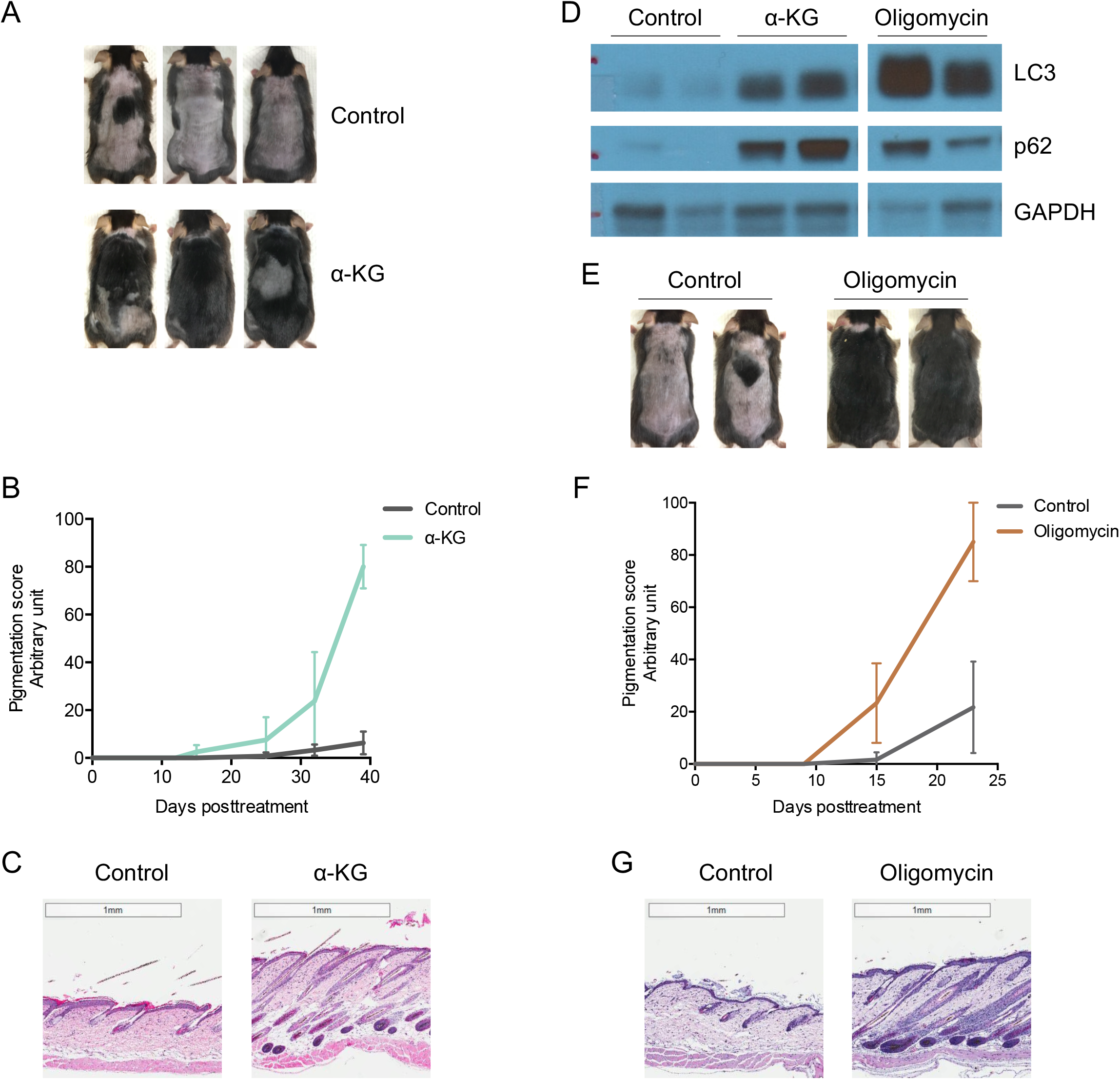
Hair regeneration is induced by topical treatment with α-KG or oligomycin. (**A**) α-KG induces hair regeneration. Mice were shaved on postnatal day 44 (telogen) and topically treated with vehicle control (DMSO) or α-KG (32 mM final in ~300 μL PLO Base) every other day over 39 days. Melanin pigmentation in the skin of α-KG treated animals, indicative of anagen induction by the treatment, became visible as early as on day 12; vehicle-treated mice did not show significant pigmentation for at least 39 days. Hair growth from the pigmented skin areas of α-KG treated mice was visible within 5~7 days. Photographs shown were taken on day 39 post-treatment, by which time mice treated with α-KG exhibited overall hair growth whereas control mice still had no hair generally except for random hair patches on some animals. Total Number of animals: control (12), α-KG (10). (**B**) Quantification for appearance of melanin pigmentation (indicating onset of anagen) in mouse skin treated with α-KG vs. control. Pigmentation scoring is described in Methods. Number of animals shown in (A): control (4), α-KG (4). (**C**) Microphotographs of hematoxylin and eosin (H&E) stained skin tissue section from mice shown in (A), showing new hair follicles, hair shafts, and thickened dermal layer in α-KG treated mouse skin. Hematoxylin is a basic dye that stains nucleic acids purplish blue; eosin is an acidic dye that stains cytoplasm and extracellular matrix (e.g., collagen) pink. (**D**) Autophagy protein LC3 is induced in telogen skin of mice treated with α-KG or oligomycin for 5 days. p62 levels are also increased. Skin remains in telogen during this whole treatment period as confirmed by the lack of skin pigmentation. Number of animals: control (3), α-KG (3), oligomycin (3). (**E**) Oligomycin (100 μM) induces hair regeneration. Photographs shown were taken on day 23 post-treatment. Number of animals: control (12), oligomycin (9). (**F**) Quantification for appearance of melanin pigmentation in mouse skin treated with oligomycin vs. control. Number of animals: control (3), oligomycin (3). (**G**) Microphotographs of H&E stained skin tissue section from mice shown in (E).

At the molecular level, longevity by α-KG was found to be mediated through direct inhibition of the highly conserved mitochondrial ATP synthase (complex V) and subsequent decrease of target of rapamycin (TOR) activity downstream (Chin et al., 2014). We tested whether hair regeneration by α-KG may also be mediated by ATP synthase inhibition. Consistent with this mechanism, the complex V inhibitor oligomycin similarly promoted hair regeneration in both male (Figure 1E-G) and female (Figure S1C) mice. Also, like α-KG, oligomycin treatment results in TOR inhibition and autophagy activation (Chin et al., 2014). Here we also detected increased autophagy in topical oligomycin treated mouse skin as indicated by increased LC3 expression (Figure 1D).

The target of rapamycin (TOR) protein is a main mediator of the effect of DR in longevity. Inhibition of TOR, e.g., by rapamycin, elicits autophagy. We therefore asked whether rapamycin may also increase hair regeneration. As shown in Figure 2A-B, topical rapamycin treatment accelerated hair regeneration, both visually and histologically. Autophagic LC3 level is increased in rapamycin treated mouse telogen skin (Figure 2C). Together, these results show that hair regeneration can be accelerated by either indirect or direct inhibition of TOR pathway activity and induction of autophagy.

**Figure 2.**
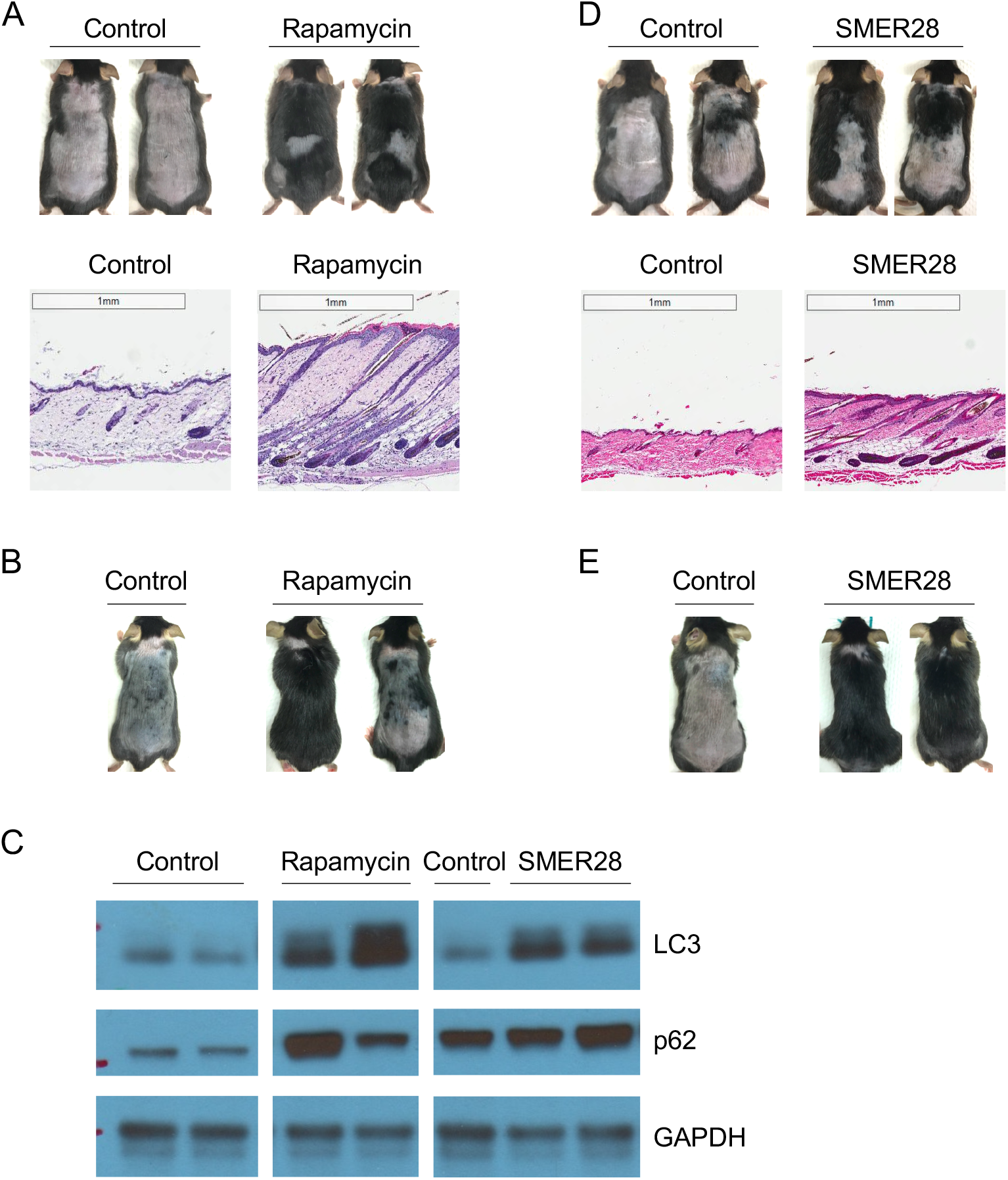
Autophagy induction is closely linked to hair regeneration. (**A**) Rapamycin (1.6 μM) induces hair regeneration. Mice were shaved on postnatal day 43 and treated topically every other day. Photographs were taken on day 41 posttreatment; H&E staining shows the entry of anagen and hair growth. Rapamycin at 16 μM does not enhance hair regeneration (data not shown); this may be due to more severe inhibition of mTOR which was reported to be required for HFSC activation (Castilho et al., 2009; Kellenberger and Tauchi, 2013; Deng et al., 2015). Number of animals: control (15), rapamycin (12). (**B**) Rapamycin (100 nM) also promotes hair regeneration. Number of animals: control (2), rapamycin (3). (**C**) LC3 and p62 in telogen skin of mice treated 5 days with indicated compounds. (**D-E**) SMER28 (1 mM) induces hair regeneration. (D) Mice were shaved on postnatal day 44 and treated every other day with topical SMER28; photographs were taken on day 23 post-treatment. Number of animals: control (6), SMER28 (5). (E) Mice were shaved on postnatal day 48 and treated daily with topical SMER28; photographs were taken on day 23 post-treatment. SMER28 has stronger effects when applied daily. Number of animals: control (1), SMER28 (3).

It was previously reported that mTOR may be required for HFSC activation and anagen entry (Castilho et al., 2009; Kellenberger and Tauchi, 2013; Deng et al., 2015). However, our results above indicate that moderate inhibition of mTOR by rapamycin accompanied by autophagy induction stimulates hair regeneration. Dichotomy may also exist for mitochondrial regulation. Mitochondrial respiration is required for HFSC cycle and genetic perturbation of mitochondrial function abolishes hair regeneration (Hamanaka et al., 2013; Shyh-Chang et al., 2013; Kloepper et al., 2015). Our finding that the well-established complex V inhibitor oligomycin in fact promotes hair regeneration suggested that possibly, as in lifespan regulation, mild mitochondrial inhibition may prove to be beneficial. Since mitochondrial complex V acts upstream of TOR from *C. elegans* to Drosophila and humans (Chin et al., 2014; Sun et al., 2014; Fu et al., 2015), and autophagy is induced both by mitochondrial complex V inhibition (Figure 1D) and by TOR inhibition (Figure 2C) (see also (Chin et al., 2014)), we further tested if autophagy induction itself is sufficient for hair regeneration. To this end, we took advantage of a TOR-independent autophagy inducing small molecule, SMER28 (Sarkar et al., 2007). Consistent with its reported function in other systems, topical SMER28 increased autophagic LC3 induction in mouse dorsal skin (Figure 2C). Importantly, SMER28 greatly induced hair regeneration (Figure 2D-E). These findings strongly support the role of autophagy, rather than mitochondrial or mTOR inhibition, in promoting hair regeneration.

Another conserved common downstream effector of α-KG and oligomycin is the AMP-activated protein kinase (AMPK) (Chin et al., 2014). AMPK is a key energy sensor of the cell. AMPK is activated by decreases in cellular energy charge, e.g., upon glucose starvation and many other cellular stress conditions (Hardie et al., 2012; Zhang et al., 2017). AMPK elevates autophagy (Galluzzi et al., 2014). Consistently, we found that, as shown in Figure 3A-C, anagen induction and hair regeneration were also stimulated by topical treatment with the AMPK activator, 5-aminoimidazole-4-carboxamide ribonucleotide (AICAR), an AMP analog. Metformin, another agonist of AMPK (Burkewitz et al., 2014), has been widely used as a diabetes drug and was the first drug approved for anti-aging human study (Barzilai et al., 2016). Its effect on hair growth has not been studied previously. Here we show that metformin similarly induced autophagy and hair regeneration (Figure 3D-G).

**Figure 3.**
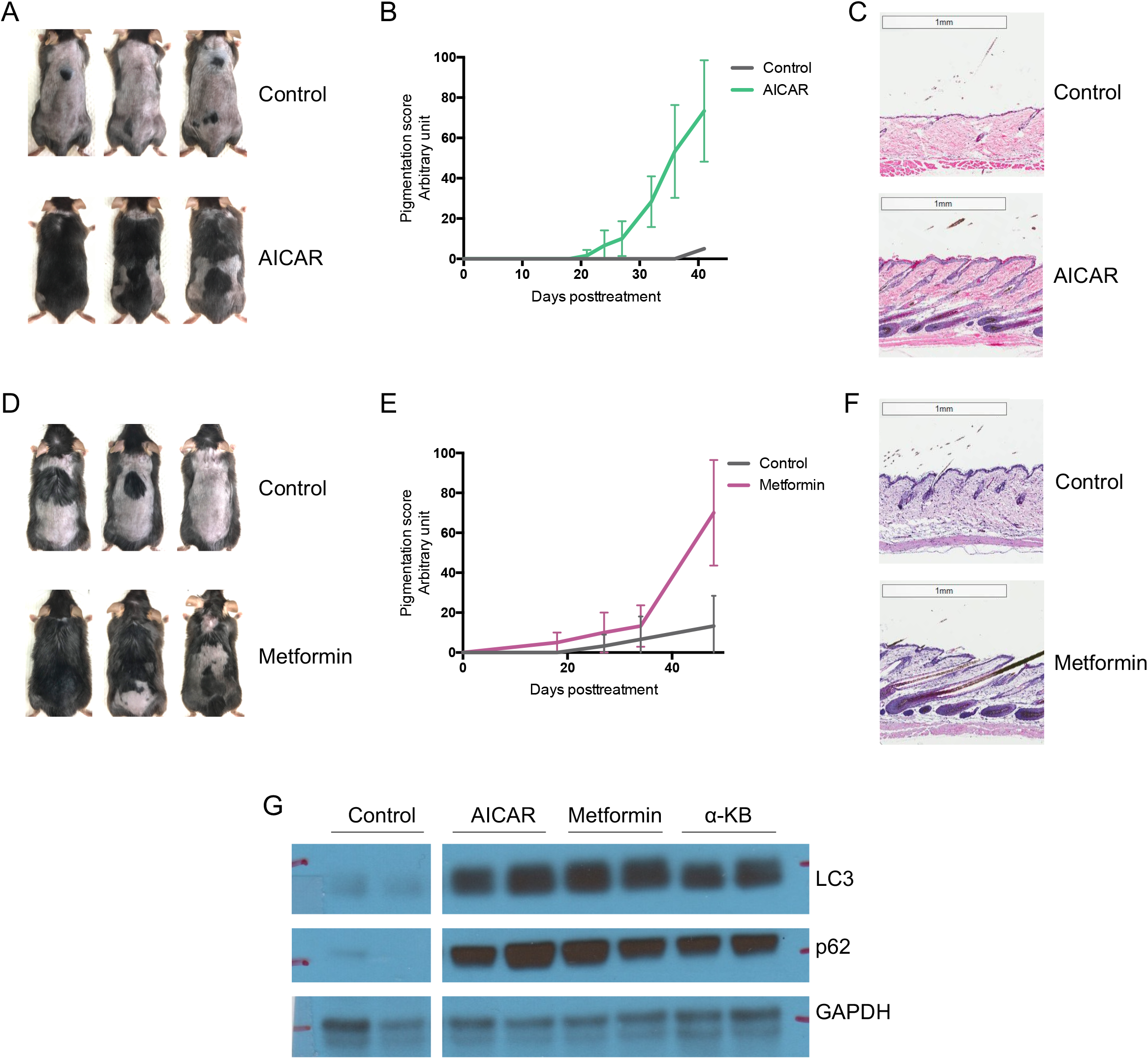
Hair regeneration is induced by AMPK activators AICAR and metformin. (**A**) AICAR (16 mM) induces hair regeneration. Mice were shaved on postnatal day 44 and treated topically every other day. Photographs were taken on day 41 post-treatment. Number of animals: control (9), AICAR (8). **(B**) Quantification for skin pigmentation in mice from (A). Number of animals: control (3), AICAR (3). (**C**) H&E stained skin tissue section from mice shown in (A). **(D**) Metformin (160 mM) induces hair regeneration. Mice were shaved on postnatal day 43 and treated topically every other day with metformin or vehicle control (H_2_O in this experiment). Photographs were taken on day 48 post-treatment. Number of animals: control (8), metformin (5). (**E**) Quantification for skin pigmentation in mice from (D). Number of animals: control (3), metformin (3). (**F**) H&E stained skin tissue section from mice shown in (D). (**G**) LC3 and p62 in telogen skin of mice treated 5 days with indicated compounds.

Mechanistically, metformin has been shown to inhibit mitochondrial complex I in the electron transport chain (Owen et al., 2000; Wheaton et al., 2014). Interestingly, long-lived *C. elegans* mitochondrial mutants accumulate various alpha keto acid metabolites in the exometabolome (Butler et al., 2010). We previously showed that one of these compounds, α-ketobutyrate (α-KB), extends the lifespan and alleviates many aging-related symptoms in the aged mice (Huang et al., 2016). α-KB supplementation in drinking water over 30 weeks greatly improved hair coating in old mice (Figure 4A). However, topical α-KB treatment in shaved aged mice only moderately increased hair regrowth, whereas α-KG or rapamycin seemed to decrease hair regeneration slightly in old mice (data not shown). On other hand, in young mice topical α-KB treatment substantially induced skin pigmentation and hair regeneration (Figure 4B-D). Furthermore, autophagy was also induced, as indicated by elevated LC3 in the treated skin (Figure 3G).

**Figure 4.**
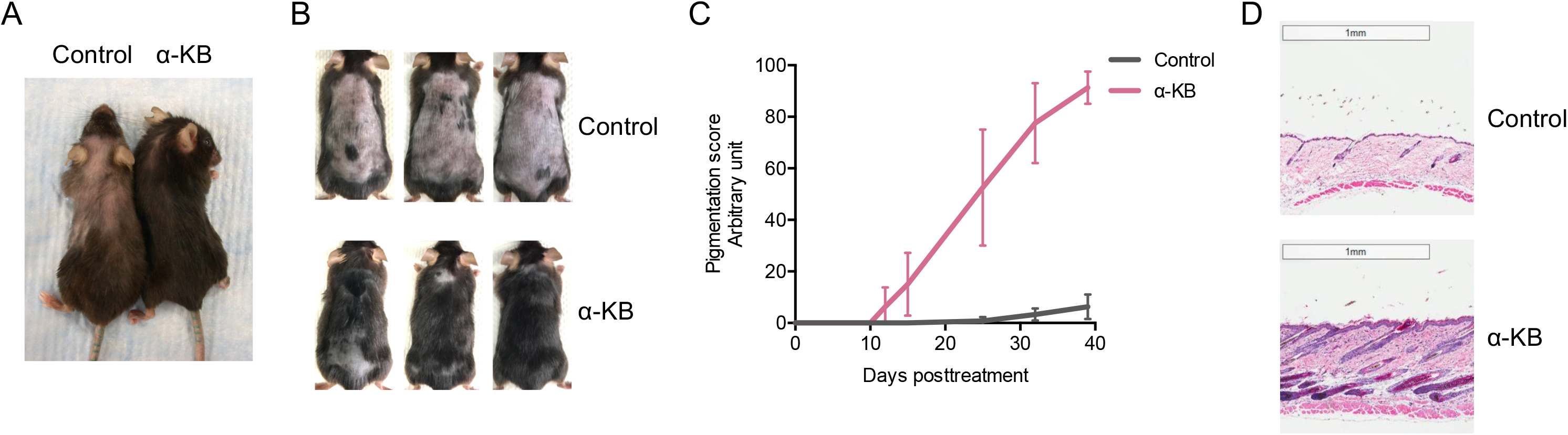
α-KB enhances hair regeneration. (**A**) Oral α-KB (8 mM in drinking water) treatment negates hair loss in aged female mice. Photo was taken at 131 weeks of age. Number of animals: control (5), α-KB (5). (**B**) Topical α-KB (32 mM) induces hair regeneration in young male mice. Mice were shaved on postnatal day 44 and treated topically every other day. Photographs were taken on day 39 post-treatment. Number of animals: control (15), α-KB (12). (**C**) Quantification for skin pigmentation in mice from (B). Number of animals: control (4), α-KB (4). (**D**) H&E stained skin tissue section from mice shown in (B).

For all molecules tested herein, autophagy induction is closely linked to hair regeneration. Moreover, we discovered that autophagy is elevated as hair follicle progresses naturally through anagen; autophagy decreases in catagen and remains low in telogen (Figure 5). These data indicate a biological role of autophagy in normal hair growth initiation and elongation. It remains to be determined whether autophagy in the epidermal, dermal, and/or hypodermal layers is entailed. In summary, we showed that autophagy induction by the longevity metabolites α-KG and α-KB, and small molecule mTOR inhibitors and AMPK activators, can initiate hair follicle activation and hair regeneration.

**Figure 5.**
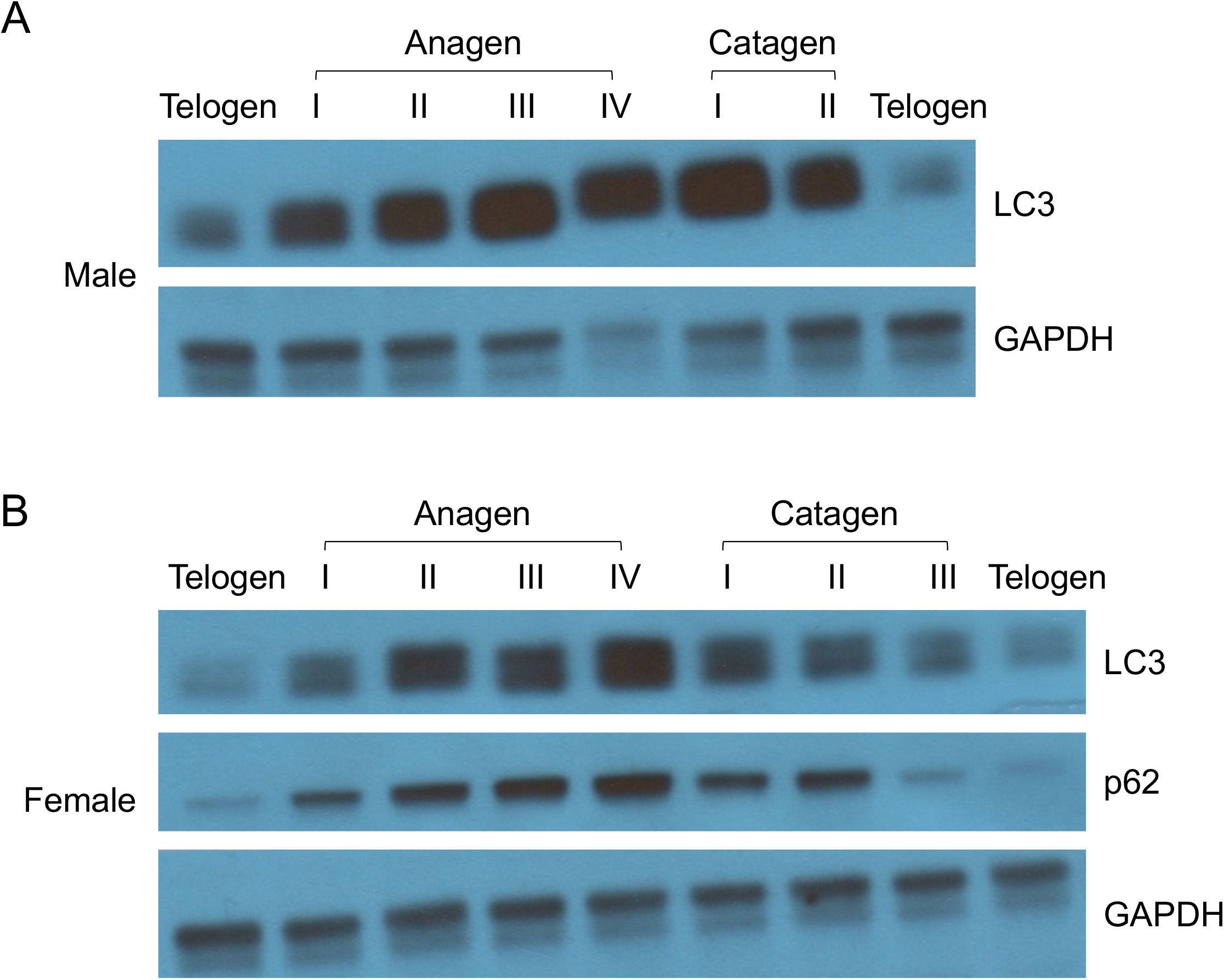
Autophagy levels are indicative of hair follicle cycle stages, increased upon anagen induction. (**A**) Male mice were shaved on postnatal day 93 and monitored for hair cycle progression. Mice at each indicated stage were sacrificed for LC3 and p62 analysis. (**B**) Female mice were shaved on postnatal day 92 and analyzed for LC3 and p62 levels at indicated hair cycle stage.

## Discussion

The regulation of hair regeneration by microenvironment, including Wnt, BMP, JAK-STAT, Treg, interleukin-2 receptor signaling, has been widely studied, but little is known about its regulation by intracellular metabolic signals (Ito et al., 2007; Harel et al., 2015; Chueh et al., 2013). Previously cellular redox (Hamanaka et al., 2013), mitochondrial integrity and energy production (Shyh-Chang et al., 2013; Kloepper et al., 2015) have been shown to be involved in HFSC activation during normal hair growth cycle which are impaired or dysregulated during aging. Here we discovered that activation of autophagy using specific small molecules can be used to promote hair regeneration.

Rapid anagen entry on whole dorsal skin was observed from time to time among mice treated with α-KG, oligomycin, rapamycin, and SMER28, but never in α-KB, metformin, or AICAR-treated mice, nor in vehicle control mice. Temporally, pigmentation (anagen entry) induction by α-KB, AICAR, or metformin takes much longer, e.g., on a time scale of 12~18 days, as compared to 5~14 days by α-KG, oligomycin, or rapamycin. It is possible that this may reflect a differential effect by mTOR inhibition from AMPK activation on the regulation of autophagy, which remains to be understood. The crosstalk between metabolism and autophagy is complex (Galluzzi et al., 2014). Autophagy is generally induced by limitations in ATP availability or a lack of essential nutrients, including glucose and amino acids. ATP is required for autophagy. Starvation and ensuing decreased energy charge and increased ROS levels are potent activators of autophagy. Recently it was reported that calorie restriction also promotes hair follicle growth and retention in mice (Forni et al., 2017) but the underlying mechanism was unclear. It is tempting to speculate that this effect of calorie restriction on hair growth may also be mediated through autophagy activation.

Autophagy has been linked to longevity but the underlying mechanisms are unclear. Although it is not sufficient for lifespan increase, many longevity pathways at least partially depend on the induction of autophagy to increase lifespan (Jia and Levine, 2007; Hansen et al., 2008). Disrupted autophagy has been linked to neurodegenerative diseases and other age-related disorders. Given the conservation in energy metabolism and autophagy machinery, induction of hair regeneration by autophagy activation discovered in mice herein should translate to humans. Although it has not been tested in human hair regeneration studies, autophagy has been shown to be essential for maintaining the growth of an ex vivo human scalp hair follicle organ culture (Parodi et al., 2018). Since α-KG is safe and has been used in dietary supplements for bodybuilding, new applications and clinical trials addressing its unprecedented anti-aging and other protective properties are highly feasible and appealing.

## Methods

### Assay for hair regeneration in mice

All compounds were tested in both male and female mice. Every experiment was repeated independently at least 2 times. Some treatments were performed concurrently with shared control arms. C57BL/6J male mice were obtained at 6 weeks of age from Jackson Laboratories (Bar Harbor, ME). C57BL/6J female mice were obtained at 8 weeks of age from Jackson Laboratories (Bar Harbor, ME). Mice were fed a standard chow diet and provided ad libitum access to food and water throughout the study. Mice were shaved dorsally in telogen, i.e., postnatal day 43~45 for males and day 58 for females, respectively. Vehicle control (25 μL DMSO, unless otherwise indicated) or test compounds (in 25 μL DMSO, unless otherwise indicated) were topically applied on the shaved skin every other day (unless otherwise described) for the duration of the experiments (4-6 weeks). Appearance of skin pigmentation and hair growth were monitored and documented by photos and videos. Progression was also assigned an arbitrary value from 0 to 100 based on skin darkening, with 0 indicating no hair growth (and no pigmentation) and higher number corresponding to darker skin and visible hair growth on larger skin areas. α-KG (Sigma 75890), oligomycin (Cell Signaling 9996L), rapamycin (Selleckchem S1039), SMER28 (Selleckchem S8240), AICAR (Selleckchem S1802), metformin (Sigma PHR1084), or α-KB (Sigma K401) in ~300 μL Premium Lecithin Organogel (PLO) Base (Transderma Pharmaceuticals Inc.) was used for each mouse.

### Aged mice

<u>For oral α-KB treatment</u>, aged male and female C57BL/6J mice were obtained at 87 weeks of age (NIA aged rodent colonies). Mice were housed in a controlled SPF facility (22 ± 2 °C, 6:00-18:00, 12 h/12 h light/dark cycle) at UCLA. Mice were fed a standard chow diet and provided ad libitum access to food and water throughout the study. Treatment with either water (vehicle control), or α-KB (90 mg/kg bodyweight) in drinking water, started when mice were at 101 weeks of age. <u>For topical α-KB treatment</u>, aged male C57BL/6J mice were obtained at 21 months of age (NIA aged rodent colonies), shaved the week after, and topically treated with α-KB (32 mM) every other day for one month. All experiments were approved by the UCLA Chancellor’s Animal Research Committee.

### Histology and microscopy

Mouse dorsal skin was shaved before being collected for histological and molecular analyses. Full-thickness skin tissue was then fixed in 10% formalin solution (Sigma, HT501128) overnight and dehydrated for embedding in paraffin. 5 μm paraffin sections were subjected to hematoxylin/eosin staining. Images were captured by Leica Aperio ScanScope AT brightfield system at X20 magnification.

### Western blotting

Male mice were shaved and treated every other day starting on postnatal day 43. After 5 days, telogen skin samples were harvested and stage confirmed. Mouse skin tissue lysate was prepared by homogenization in T-PER Tissue Protein Extraction Buffer (Thermo Scientific, 78510) with protease inhibitors (Roche, 11836153001) and phosphatase inhibitors (Sigma, P5726) by FastPrep-24 (MP Biomedicals). Tissue and cell debris was removed by centrifugation and the lysate was boiled for 5 min in 1 x SDS loading buffer containing 5% β-mercaptoethanol. Samples were then subjected to SDS-PAGE on NuPAGE Novex 4-12% Bis-Tris gradient gels (Invitrogen, NP0322B0X), and western blotting was carried out with antibodies against LC3 (Novus, NB100-2220), p62 (Sigma, P0068), or GAPDH (Ambion, AM4300).

### Statistical Analysis

All treatments were repeated at least two times with identical or similar results. Data represent biological replicates. Appropriate statistical tests were used for every figure. Data meet the assumptions of the statistical tests described for each figure. Mean ± s.d. is plotted in all figures unless stated otherwise.

## Acknowledgment

We thank the Translational Pathology Core Laboratory at UCLA for core services, H. Hwang, B. Lomenick, W. Lowry, and M. Miranda for technical advice. This work was partially supported by the Margaret E. Early Medical Research Trust, LongLifeRx Inc, and NIH grants P01 HL090553 and AG049753.

## Supplemental Information

**Figure S1.**
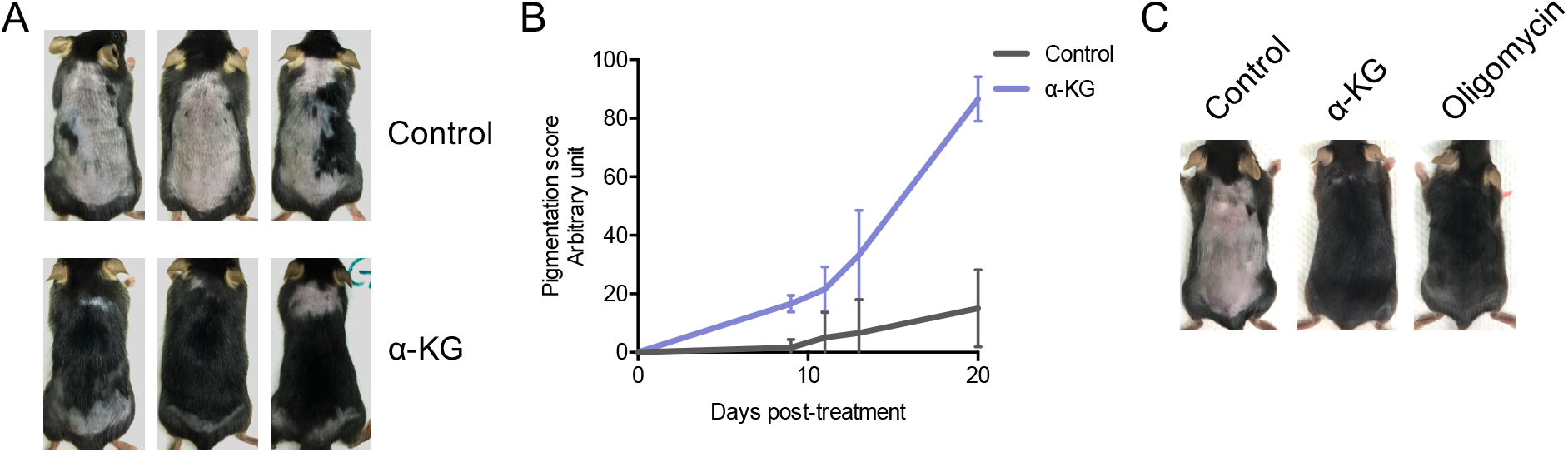
In both male and female mice, hair regeneration can be induced by α-KG or oligomycin treatment. (**A**) Compared to 6.5-week old mice in Figure 1A, α-KG induces faster hair regeneration in 8-week old animals. Male mice were shaved on postnatal day 57 (telogen) and topically treated every other day. Photographs shown were taken on day 20 posttreatment. Number of animals: control (4), α-KG (3). (**B**) Quantification for skin pigmentation in mice from Figure S1A. Pigmentation of α-KG treated animals became visible as early as on day 7, and full dorsal hair coverage was observed by day 20 post-treatment. Number of animals: control (3), α-KG (3). (**C**) α-KG and oligomycin also stimulate hair regeneration in female animals. Female mice were shaved on postnatal day 58 (telogen) and topically treated with vehicle control (DMSO), α-KG, or oligomycin every other day. Photographs shown were taken on day 26 post-treatment. Number of animals: control (4), α-KG (2), oligomycin (4).

